# SqueezeCall: Nanopore basecalling using a Squeezeformer network

**DOI:** 10.1101/2025.01.21.634194

**Authors:** Zhongxu Zhu

## Abstract

Nanopore sequencing, a novel third-generation sequencing technique, offers significant advantages over other sequencing approaches, owing especially to its capabilities for direct RNA sequencing, real-time analysis, and long-read length. During nanopore sequencing, the sequencer measures changes in electrical current that occur as each nucleotide passes through the nanopores. A basecaller identifies the base sequences according to the raw current measurements. However, due to variations in DNA and RNA molecules, noise from the sequencing process, and limitations in existing methodology, accurate basecalling remains a challenge. In this paper, we introduce SqueezeCall, a novel approach that uses an end-to-end Squeezeformer-based model for accurate nanopore basecalling. In SqueezeCall, convolution layers are used to down sample raw signals and to model local dependencies. A Squeezeformer network is employed to capture the global context. Finally, a connectionist temporal classification (CTC) decoder generates the DNA sequence by a beam search algorithm. Inspired by the Wav2vec2.0 model, we masked a proportion of the time steps of the convolution outputs before feeding them to the Squeezeformer network and replaced them with a trained feature vector shared between all masked time steps. Experimental results demonstrate that this method enhances our model’s ability to resist noise and allows for improved basecalling accuracy. We trained SqueezeCall using a combination of three types of loss: CTC-CRF loss, intermediate CTC-CRF loss, and KL loss. Ablation experiments show that all three types of loss contribute to basecalling accuracy. Experiments on multiple species further demonstrate the potential of the Squeezeformer-based model to improve basecalling accuracy and its superiority over a recurrent neural network (RNN)-based model and Transformer-based models.

## INTRODUCTION

Nanopore sequencing, a novel third-generation sequencing technique, has undergone substantial improvement in the past several years [1, 2]. These advances have highlighted key advantages of nanopore sequencing, particularly its capabilities for high-throughput and real-time analysis, long-read lengths, and direct RNA sequencing. Nanopore sequencing works by measuring changes in electric current that are generated by the movement of DNA or RNA molecules through nanopores. Basecalling is usually the initial step in analyzing nanopore sequencing signals. The basecaller deduces base sequences from raw current measurements. Nearly all downstream applications depend on this fundamental basecalling process.

Basecalling is a challenging task for several reasons. First, a nanopore can hold *k* nucleotides (*k*-mer) simultaneously (e.g., 5 for the pore version R9.4). Thus, the current signal level does not correspond to a single base but is most dominantly influenced by the *k*-mers that pass through the nanopores. Second, the inherent sequencing limitation of nanopore technology leads to low signal-to-noise ratio of the original raw data, which makes it difficult to identify the true nucleotide sequence.

The early development of basecallers can be divided roughly into two stages. In the first stage, current measurements were converted to events, each corresponding to the movement of a *k*-mer through the pore [1]. Then a hidden Markov model (HMM) and a Viterbi decoding algorithm was implemented to model the event space and further decode the base sequence. In the second stage, end-to-end deep learning-based approaches became popular for basecalling; these approaches generate the DNA sequence directly from raw electrical data. Chiron [2] used developments in the automatic speech recognition (ASR) field as it applied a convolutional neural network (CNN) to extract features from the raw signal, to a recurrent neural network (RNN) to relate such features in a temporal manner, and to a connectionist temporal classification (CTC) decoder [3] to generate a variant-length base sequence for a fixed-length signal window through output-space searching. Mincall [4] used a deep CNN with residual connections [5], batch normalization, and CTC loss. Causalcall [6] used a modified temporal convolutional network (TCN) [17] with causal dilations [7] and a CTC decoder.

In contrast to the basecalling models using RNNs, the convolution-based model of Causalcall can speed up basecalling by matrix computation. SACall [8] used convolution layers to downsample the signals and capture local patterns and adopted self-attention layers [10] to calculate the similarity of the signals at any two positions in the raw signal sequence. It then used the CTC decoder to generate the DNA sequence by a beam search algorithm. Other approaches have included CATCaller [9], which employed lite-transformers [11] to capture global context; URNano [12], which utilized a convolutional U-net with integrated RNNs [13]; and Halcyon [14], which used a sequence-to-sequence (Seq2Seq) model with attention [15][16]. The official tool of Oxford Nanopore Technologies (ONT) is Bonito [30], a CNN- and LSTM-based tool that captures both local and global patterns and employs a conditional random field (CRF) decoder to generate a base sequence. Recently Bonito updated a transformer version with the capability to learn global patterns.

CNN and Transformer architectures are widely used in the basecall field and currently are the popular backbone architecture for ASR models. But they have limitations. In general, the CNN model lacks the ability to capture the global context, and Transformer is expensive in terms of computation and memory. To overcome these shortcomings, Conformer, a new convolution-augmented Transformer architecture, was developed. Conformer captures global and local features synchronously from audio signals. Squeezeformer is a further improvement upon Conformer. Conformer was found [20] to have a high temporal redundancy in the learned feature representations of neighboring speech frames, which results in unnecessary computational overhead. To address this, Squeezeformer incorporated the temporal U-Net structure, in which a downsampling layer halves the sampling rate at the middle of the network and a light upsampling layer recovers the temporal resolution at the end for training stability. Squeezeformer achieved a better word-error-rate (WER) on LibriSpeech with the same amount of computation.

In this paper, we propose an end-to-end Squeezeformer-based model for accurate nanopore basecalling named SqueezeCall. The convolution layers are used to downsample the raw signals and model the local dependencies, and a Squeezeformer network is employed to capture global context. Inspired by the Wav2vec2.0 model, we added a mask module between the convolution network and the Squeezeformer network. This addition masked a proportion of the time steps of the convolution outputs before feeding them to the Squeezeformer network and replaced them with a trained feature vector shared between all masked time steps. Experimental results demonstrate that this method increases our model resistance to noise and thus improves accuracy. We used a combination of three types of loss: CTC-CRF loss[28][29][30], intermediate CTC-CRF loss [23], and KL loss in the training process. Ablation study demonstrates that each loss contributes significantly to the accuracy rate. We compared SqueezeCall with four basecaller methods, including Bonito-LSTM, Bonito-Transformer, CATCaller, and SACall, on 11 datasets (NA12878 Human Dataset, Lambda Phage, and nine bacterial datasets).

## MATERIALS AND METHODS

### A. Datasets

We used a human genome reference (NA12878/GM12878, Ceph/Utah pedigree) dataset from [25]. The human dataset contained many different sequencing runs. We chose three experiments: FAB42828, FAF09968, and FAF04090. We adopted the same train-test-split method as in [27], training on approximately 80,000 reads and testing on 5,000 reads. We also included a Lambda phage dataset [27], which was trained on approximately 40,000 reads and tested on 5,000 reads.

We also used the bacterial dataset released by Wick et al. [26]. This dataset contains two parts: training dataset and test dataset. The training dataset consists of 50 individual species genomes, consisting of 30 *Klebsiella pneumoniae* genomes, 10 genomes of other species of Enterobacteriaceae, and 10 genomes from other families of Proteobacteria. The test dataset is composed of nine species, including three *Klebsiella pneumoniae, Shigella sonnei, Serratia marcescens, Haemophilus haemolyticus, Acinetobacter pittii, Stenotrophomonas maltophilia*, and *Staphylococcus aureus*.

To obtain accurate label sequences and increase the quality and usability of our training data, data was annotated using the Tombo resquiggle tool (v1.5.1). First, reads were aligned to the reference sequences using their basecalls. Reference genomes were used as reference sequences for all the datasets. Using Tombo, we aligned the raw signal to the expected signal according to the reference. Reads that did not align to the reference genome or that provided a bad resquiggle quality according to Tombo were discarded.

In addition, we used the public dataset for ablation studies provided by ONT [18]. The dataset divides reads into chunks of equal length and consists of a training set of 1.22 million chunks and a development set of 20,000 chunks, each with 3,600 current signal values.

### B. Squeezeformer architecture

The structure of Squeezeformer is shown in Figure 1, which illustrates the Depthwise Separable Conv Subsampling module and the Temporal U-Net Architecture of the encoder. The Depthwise Separable Conv Subsampling module subsamples the original current signals with a sampling rate of *R*_0_. The Temporal U-Net Architecture network consists of 2N Squeezeformer blocks. The first N-1 Squeezeformer blocks maintain the same sampling rate of *R*_0_ and then enter the middle N blocks after 2x downsampling through the pooling layer. The middle N Squeezeformer blocks maintain the same sampling rate of *R*_0_/2 and are connected with residual. The last Squeezeformer block is upsampled, and the sampling rate returns to R0. The Conformer module consists of a feed-forward (F) network, multi-head attention (MHA), Convolution (C), and another feed-forward (F) network; this module is referred to as the FMCF structure.

**Figure 1.**
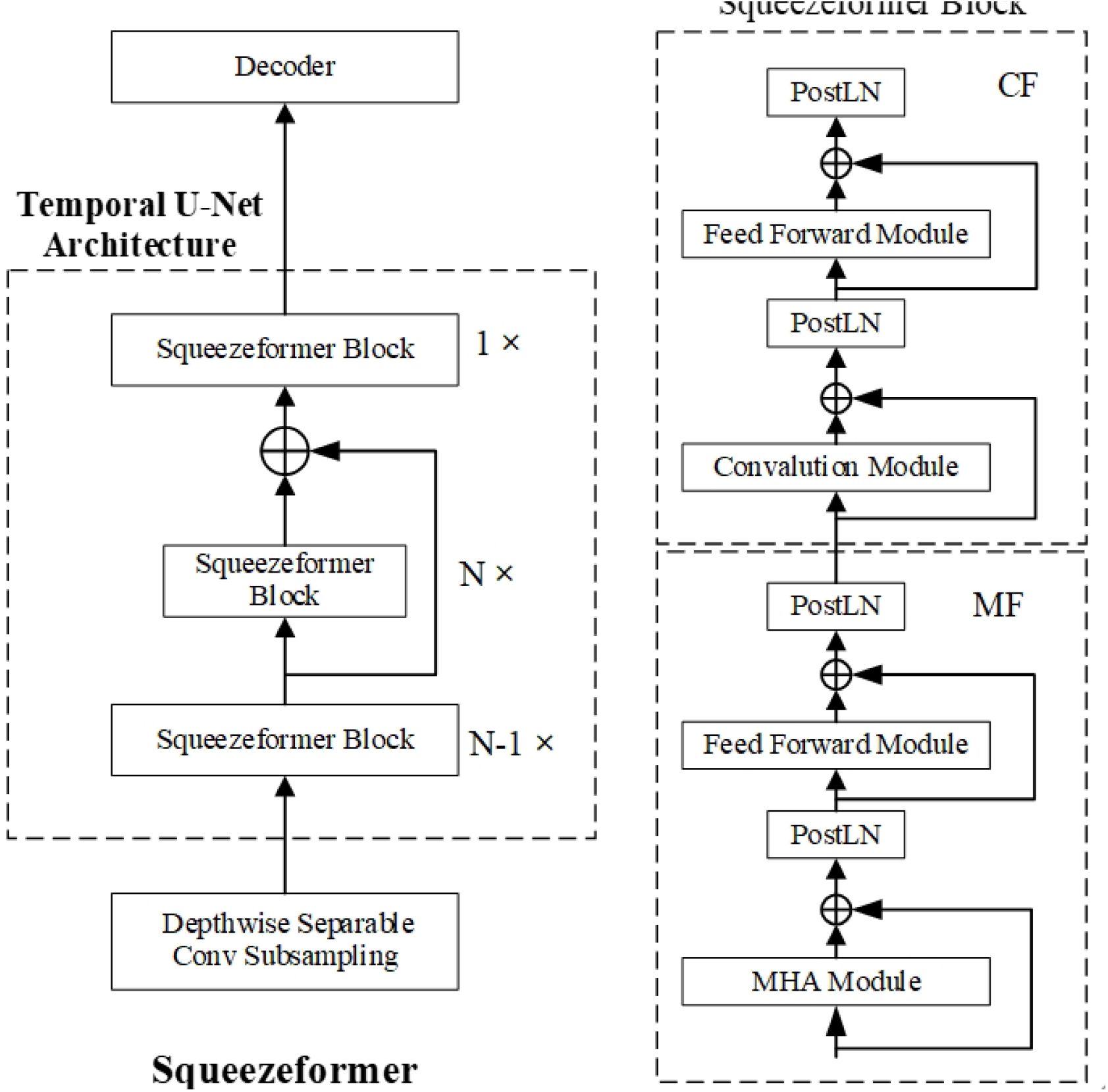
The Squeezeformer architecture consists of the Temporal U-Net structure for downsampling and upsampling of the sampling rate, the standard Transformer-style block structure that only uses Post-Layer Normalization, and the depthwise separable subsampling layer.

### C. SqueezeCall Model architecture

The structure of the SqueezeCall model is shown in Figure 2. The encoder section consists of three modules:

**Figure 2.**
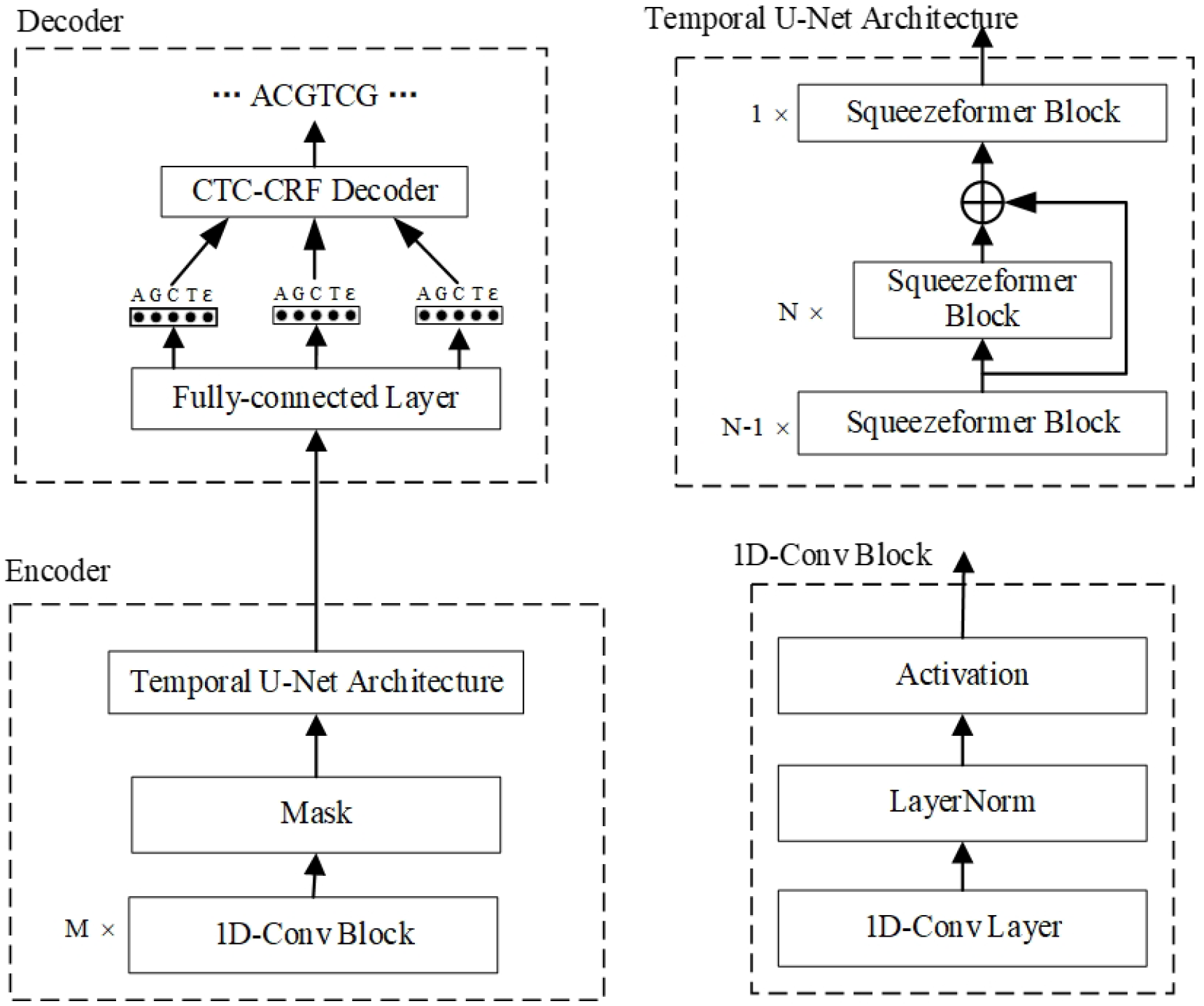
The structure of SqueezeCall

#### Convolution module

The convolution module contains two 1D convolution blocks, each of which consists of a normal 1D convolution layer followed by a layer normalization and a Gaussian Error Linear Unit (GeLU) function [22]. The first and second convolution layer have a kernel = 5, padding = 1, and stride = 1. The third convolution layer has a kernel = 19, padding = 10, and stride = 1. The output channels are 4, 16, and 512 respectively.

#### Mask module

Inspired by the Wav2vec2.0 model [21], we added a max module between the Convolution module and the Temporal U-Net Architecture. We first masked a proportion of the feature encoder outputs (time steps) before feeding them to the Temporal U-Net Architecture and replaced them with a trained feature vector shared between all masked time steps. A certain proportion of all time steps were randomly sampled without replacement as starting indices; we then masked the subsequent *mask_time_length* consecutive time steps from every sampled index. We also tried to mask in the feature dimension, but there was no significant improvement in accuracy.

#### Temporal U-Net Architecture

We employ the Temporal U-Net Architecture in Squeezeformer to capture global context. The Temporal U-Net Architecture network consists of 2N Squeezeformer blocks. The first N-1 Squeezeformer blocks maintain the same sampling rate of *R*_0_ and then enter the middle N blocks after 2x downsampling through the pooling layer. The middle N Squeezeformer blocks maintain the same sampling rate of *R*_0_/2, and the last one is upsampled, with the sampling rate returning to R_0_.

We employed a CTC decoder to generate the base sequence. After the encoder module is a fully connected layer followed by a log-softmax function to convert hidden states at position *j* to log probabilities:

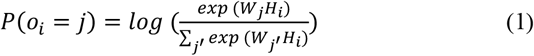

The output *o*_*i*_ predicts the corresponding symbol *j* = {A, G, C, T, ε}, where ε stands for a blank symbol. The output is then fed into a CTC decoder based on a beam search algorithm with a beam width w = 5. The beam search decoder maintains a set of prefix sequences with the maximum probability of 0∼i positions. The new set of prefixes at the i+1 position is generated on top of the previous set of prefixes, each extending all possible characters and preserving the top-ranked candidate prefix. The algorithm accumulates the scores of each prefix in the beam in each iteration of the search process and ultimately selects the one with the highest score as the final output sequence.

### D. Object

We used a combination of three types of loss: CTC-CRF loss[28][29][30], intermediate CTC-CRF loss, and KL loss in the training process.

#### Intermediate CTC-CRF loss

The intermediate CTC-CRF loss in *l-*th block is defined as:

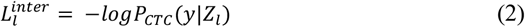

Where *Z*_*l*_ represents the output of the *l*-th block, and *y* is the label sequence. The final intermediate CTC-CRF loss is the sum of intermediate loss of multiple intermediate blocks, which is defined as follows:

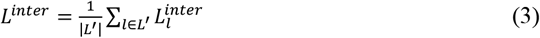

where *L*^′^ represents the block set for which intermediate loss is selected, which can be multiple blocks or a single block.

#### KL loss

The KL loss measures the Kullback-Leibler divergence between the predicted probability distribution and the true distribution, which is defined as follows:

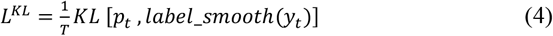

where *p*_*t*_ represents the probability distribution at time *t*, and *y*_*t*_ represents the label value at time *t*. The factor *label_smooth* (*) indicates that the Label Smoothing function and smooth factor is 0.1. For example, if the label set is five classes of {A, G, C, T, ε}, *y*_*t*_ *=* 1, *label_smooth*(*y*_*t*_) is 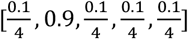. We selected only the N-th block for intermediate CTC-CRF loss (total of 2N blocks).

The final loss is a combination of three types of loss:

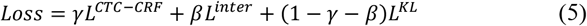

where γand β are the weight coefficients. We found that γ = 0.3 and β = 0.35 achieved good results.

### E. Training setting

Reads were sliced in non-overlapping chunks of 3,600 data points for training efficiency. We use Adam [24] to optimize the above loss function. The learning rate is warmed up over the first 1,000 updates to a peak value of 0.0005 and then linearly decayed. We trained on 4 GPUs with a batch size of 16 per GPU, giving a total batch size of 64.

## RESULTS AND DISCUSSION

### Performance comparison

We constructed our model variants with different sizes: SqueezeCall-M and SqueezeCall-L. They have 8 and 10 layers and 79M and 95M parameters, respectively. We compared SqueezeCall with four basecaller methods, including Bonito-LSTM, Bonito-Transformer, CATCaller, and SACall, on 11 datasets (NA12878 Human Dataset, Lambda Phage, and nine bacterial datasets). Bonito-LSTM and Bonito-Transformer are ONT official state-of-the-art basecall tools, which have 27M and 79M parameters, respectively.

### Error Rate of Basecalled Reads

Table 1 presents the error rate of six basecaller methods, including SqueezeCall-L, SqueezeCall-M, Bonito-LSTM, Bonito-Transformer, CATCaller, and SACall. Deletion, insertion, and mismatch rates are defined as the number of deleted, inserted, and mismatched bases divided by the alignment length. Error rate is defined as the sum of deletion, insertion, and mismatch rates. It should be noted that the error rate of CATCaller and SACall on the human dataset came from [27], and the error rate of SACall on the bacterial datasets came from [8]. CATCaller did not report the detailed error rate on the bacterial datasets. SqueezeCall-L achieved the lowest error rate across all datasets, followed by SqueezeCall-M. Despite the Bonito-Transformer achieving the lowest insertion rate and mismatch rate on some datasets, its overall error rate is higher than that of SqueezeCall.

**Table 1.**
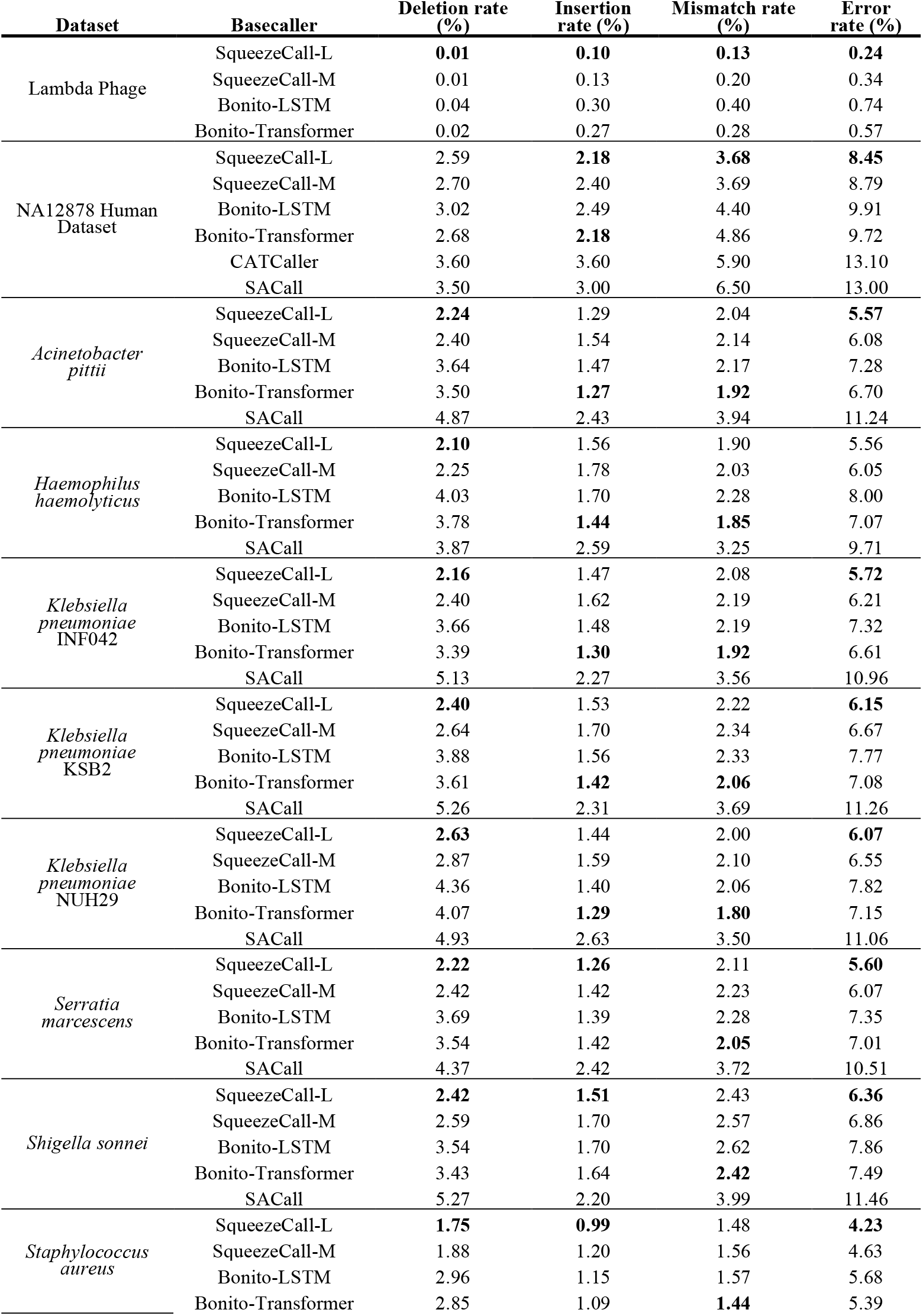
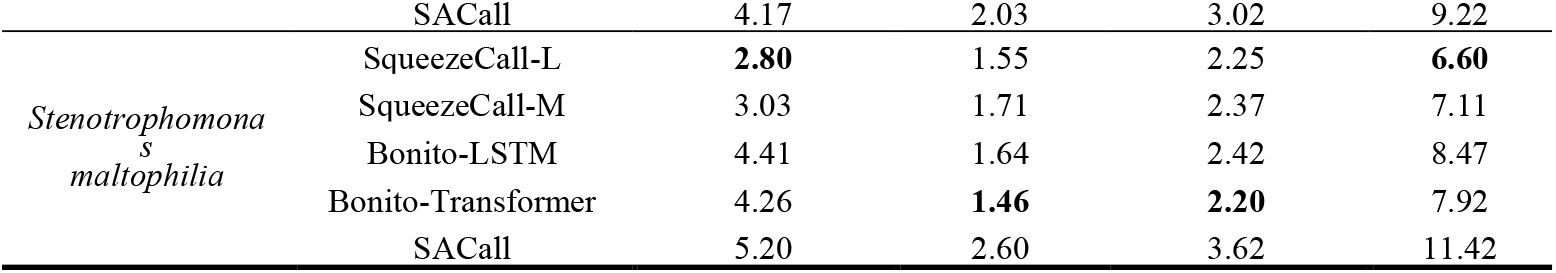
Read Level Error Rate of Three Basecalling Models on the Test Dataset.

Homopolymer and heteropolymer regions are the key challenges in nanopore sequencing. A detailed analysis of homopolymer and heteropolymer regions were performed. As is showed in Table 2 the error rate rises rapidly as the length increases. The error rate was higher for C/G homopolymer compared to A/T homopolymer while high probability of skipping was observed (Figure 3). Compared with bonitoLSTM and bonitoTransformer, SqueezeCall had a higher accuracy in homopolymer and heteropolymer regions (Figure 3 and Table 2).

**Table 2.** Accuracy of homopolymer and heteropolymer.

| Length | Homopolymer Accuracy |  |  | Heteropolymer Accuracy* |  |  |
| --- | --- | --- | --- | --- | --- | --- |
|  | SqueezeCall | bonitoLSTM | bonitoTransformer | SqueezeCall | bonitoLSTM | bonitoTransformer |
| 3 | **0.90** | 0.89 | 0.89 |  |  |  |
| 4 | **0.85** | 0.81 | 0.83 | **0.91** | 0.87 | 0.88 |
| 5 | **0.76** | 0.68 | 0.72 |  |  |  |
| 6 | **0.66** | 0.57 | 0.64 | **0.84** | 0.81 | 0.81 |
| 7 | **0.57** | 0.49 | 0.50 |  |  |  |
| 8 | **0.32** | 0.29 | 0.27 | **0.78** | 0.68 | 0.70 |
| 9 | 0.07 | 0.09 | **0.10** |  |  |  |
| 10 |  |  |  | **0.86** | 0.76 | 0.77 |

\* Heteropolymer refers to repeated dimers.

**Figure 3.**
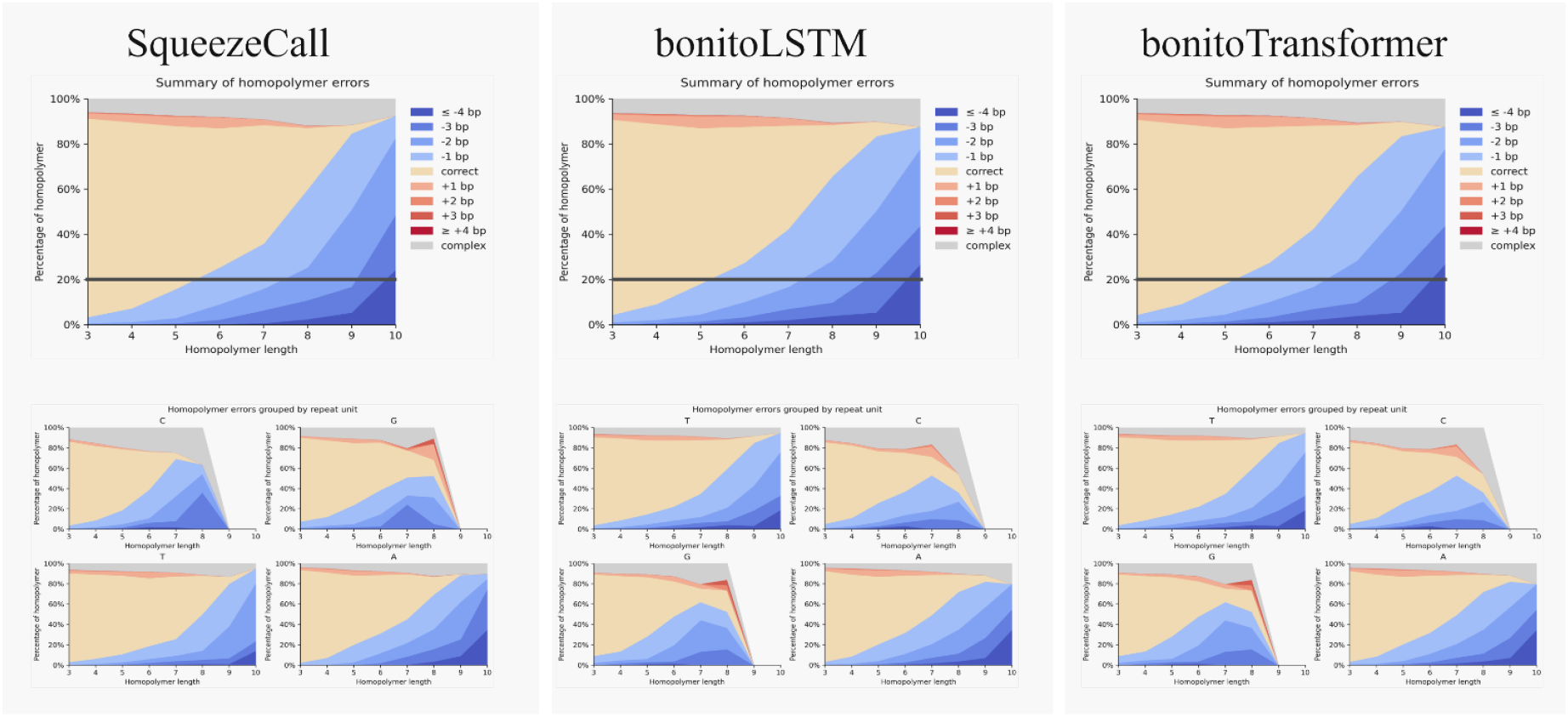
The homopolymer errors grouped by repeat unit

### Read Identity of Basecalled Reads

Figure 4 presents the identity rate of the six basecaller methods. The identity rate is defined as the portion of the read length that aligns correctly. SqueezeCall-L achieved the highest identity rate, at 93.97%, on average^1^. Average identity rates for SqueezeCall-M, Bonito-Transformer, Bonito-LSTM, CATCaller, and SACall were 93.50, 92.79, 92.25, 91.2, and 90.66%, respectively.

**Figure 4.**
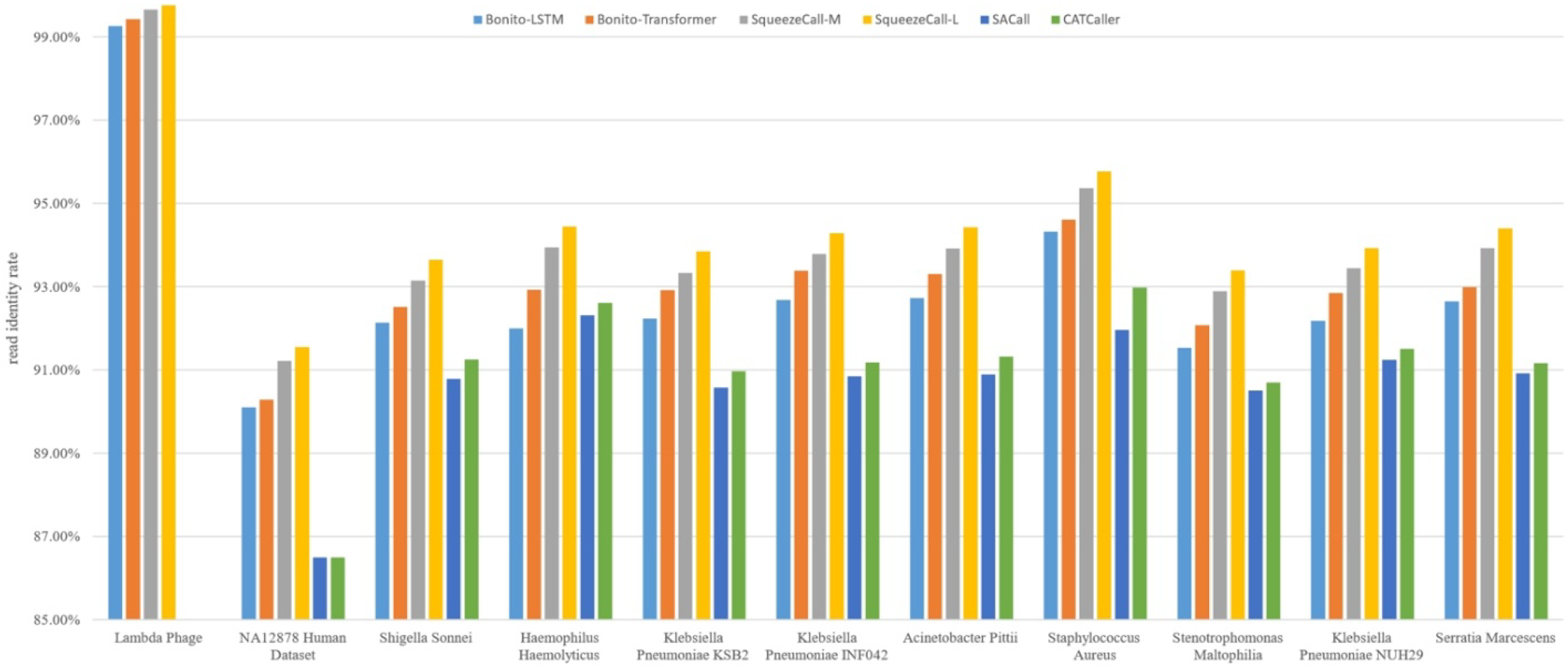
The read level identity rate of six basecallers on the 11 datasets.

### Ablation studies

The results of our ablation experiments are reported in terms of the median match rate of data chunks.

### Convolution module

Table 3 shows the results of ablation studies using the convolution module. After removing LayerNorm from SqueezeCall-M, the median match rate decreased by 0.17%. This demonstrated that adding a LayerNorm after a convolution layer can help to capture local patterns. Next, we increased the number of convolution layers to 4 and 5. However, we found no further improvement in terms of median match rate, suggesting that the 3-layer convolution networks are sufficient to learn local information.

**Table 3.**
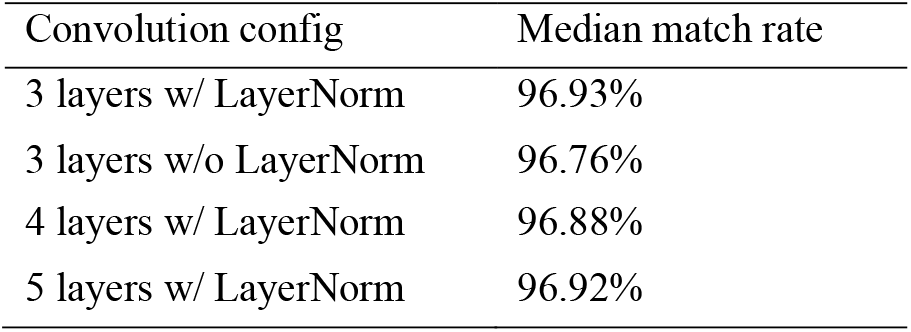
Ablation studies of the convolution module.

| Convolution config | Median match rate |
| --- | --- |
| 3 layers w/ LayerNorm | 96.93% |
| 3 layers w/o LayerNorm | 96.76% |
| 4 layers w/ LayerNorm | 96.88% |
| 5 layers w/ LayerNorm | 96.92% |

### Mask module

We experimented with various values for ‘mask_time_prob’ and ‘mask_time_length’, including ‘mask_time_prob’ values of 0.01, 0.05, and 0.1, and ‘mask_time_length’ values of 3, 5, and 10 (Table 4). The experimental results indicate that the highest Median match rate, at 96.93%, was achieved when ‘mask_time_prob’ was set to 0.05 and ‘mask_time_length’ was set to 5. Conversely, the lowest Median match rate, at 96.21%, was obtained when ‘mask_time_prob’ was set to 0.1 and ‘mask_time_length’ was set to 3. After removing the mask module from the SqueezeCall-M model, we found that the median match rate decreased from 96.93% to 96.76%. We suspect that the max module increases our model resistance to noise, thereby improving accuracy.

**Table 4.** The selection of hyperparameters for *mask_time_prob* and *mask_time_length*.

| mask_time_prob | mask_time_length | Median match rate |
| --- | --- | --- |
| 0.01 | 3 | 96.53% |
| 0.01 | 5 | 96.72% |
| 0.01 | 10 | 96.29% |
| 0.05 | 3 | 96.75% |
| 0.05 | 5 | 96.93% |
| 0.05 | 10 | 96.90% |
| 0.1 | 3 | 96.21% |
| 0.1 | 5 | 96.31% |
| 0.1 | 10 | 96.23% |

### Combination of three losses

We used the SqueezeCall-M model to study the effects of three losses: CTC-CRF loss, Intermediate CTC-CRF loss, and KL loss. Table 5 shows the impact of adding one loss in turn. Using only CTC-CRF loss, we achieved a 96.37% median match rate. Combining the CTC-CRF loss and the intermediate loss with weights 0.55/0.45 resulted in a significant performance improvement (96.37% vs 96.64%). The result indicates that intermediate loss relaxes the conditional independence assumption of CTC-based basecalling, helping to improve accuracy. Finally, we combined three kinds of losses with weights 0.35, 0.30, and 0.35 and found that the median match rate is further reduced by 0.29% (96.64% vs 96.93%).

**Table 5.** Ablation studies of three kinds of loss. *(1) CTC-CRF loss; (2) adding intermediate loss; and (3) adding KL loss*.

| Loss change | Median match rate |
| --- | --- |
| CTC-CRF loss | 96.37% |
| + Intermediate CTC-CRF loss | 96.64% |
| + KL loss | 96.93% |

## CONCLUSIONS

The accuracy of nanopore sequencing still needs to be improved. With the advancement of deep learning, we introduce SqueezeCall, a novel end-to-end Squeezeformer-based model for accurate basecalling in this paper. SqueezeCall uses convolution layers to downsample raw signals, a Squeezeformer network to capture global context, and a CTC decoder to generate the DNA sequence. Inspired by the Wav2vec2.0 model, we masked a proportion of the time steps of the convolution outputs before feeding them to the Squeezeformer network. We then replaced them with a trained feature vector shared between all masked time steps. This approach both increases resistance to noise and improves accuracy. We further demonstrate, through ablation studies, that different types of loss contribute significantly to the accuracy rate. We compared SqueezeCall with four established basecaller methods, and found that SqueezeCall achieved the highest read identity rate. SqueezeCall can be integrated into upstream analysis of sequencers to improve the accuracy of basecalling which could facilitate downstream genome assembly and environmental sample analysis. Furthermore, SqueezeCall holds the potential to directly call modified bases. In future studies, training on highly curated datasets including known modifications will increase research value.

## AVAILABILITY OF SUPPORTING SOURCE CODE AND REQUIREMENTS

- Project name: SqueezeCall
- Project home page: https://github.com/labcbb/SqueezeCall
- Operating system(s): Platform independent
- Programming language: Python
- License: MIT
- biotools ID: squeezecall
- RRID: SCR_026321

## ACKNOWLEDGEMENTS

Not applicable.

## DATA AVAILABILITY

Human raw data and basecall datasets (FAF04090, FAF09968, FAB42828) are available at https://github.com/nanopore-wgs-consortium/NA12878/blob/master/Genome.md [25]; bacterial raw data and lambda phage data are available at https://github.com/marcpaga/nanopore_benchmark/tree/main/download [26][27]; ONT chunk dataset are available at https://cdn.oxfordnanoportal.com/software/analysis/bonito/example_data_dna_r9.4.1_v0.zip [18].

## DECLARATIONS

### Ethics approval and consent to participate

Not applicable.

### Authors’ contributions

ZZ, conceptualization, project administration, software, data collection, processing, and application, manuscript writing and figure.

## Footnotes

1 Since the identity rate of the Lambda Phage dataset is significantly higher than that of other datasets, the Lambda Phage data was not included when calculating the average identity rate.

